# A novel assay to isolate and quantify third-stage filarial larvae emerging from individual mosquitoes

**DOI:** 10.1101/2020.02.04.934653

**Authors:** Abigail R. McCrea, Elizabeth B. Edgerton, Genevieve T. Oliver, Fiona M. O’Neill, Thomas J. Nolan, James B. Lok, Michael Povelones

## Abstract

**Background:** Mosquitoes transmit filarial nematodes to both human and animal hosts, resulting in worldwide health and economic consequences. Transmission to a vertebrate host requires that ingested microfilariae develop into infective third-stage larvae capable of emerging from the mosquito proboscis onto the skin of the host during blood feeding. Determining the number of microfilariae that successfully develop to infective third-stage larvae in the mosquito host is key to understanding parasite transmission potential and to developing new strategies to block these worms in their vector.

**Methods:** We developed a novel method to efficiently assess the number of infective third-stage filarial larvae that emerge from experimentally infected mosquitoes. Following infection, individual mosquitoes were placed in wells of a multi-well culture plate and warmed to 37 °C to stimulate parasite emergence. *Aedes aegypti* infected with *Dirofilaria immitis* were used to determine infection conditions and assay timing. The assay was also tested with *Brugia malayi* infected *Ae. aegypti*.

**Results:** Approximately 30% of *Ae. aegypti* infected with *D. immitis* and 50% of those infected with *B. malayi* produce emerging third-stage larvae. Once *D. immitis* third-stage larvae emerge at 13 days post infection, the proportion of mosquitoes producing them, and the number produced per mosquito remain stable until at least day 21. The prevalence and intensity of emerging third-stage *B. malayi* were similar on days 12-14 days post infection. Increased uptake of *D. immitis* microfilariae increases the fitness cost to the mosquito but does not increase the number of emerging third-stage larvae.

**Conclusions:** We provide a new assay with an associated set of infection conditions that will facilitate assessment of the filarial transmission potential of mosquito vectors and promote preparation of uniformly infectious L3 for functional assays. The ability to quantify infection outcome will facilitate analyses of molecular interactions between vectors and filariae, ultimately allowing for the establishment of novel methods to block disease transmission.

**Graphical Abstract:** 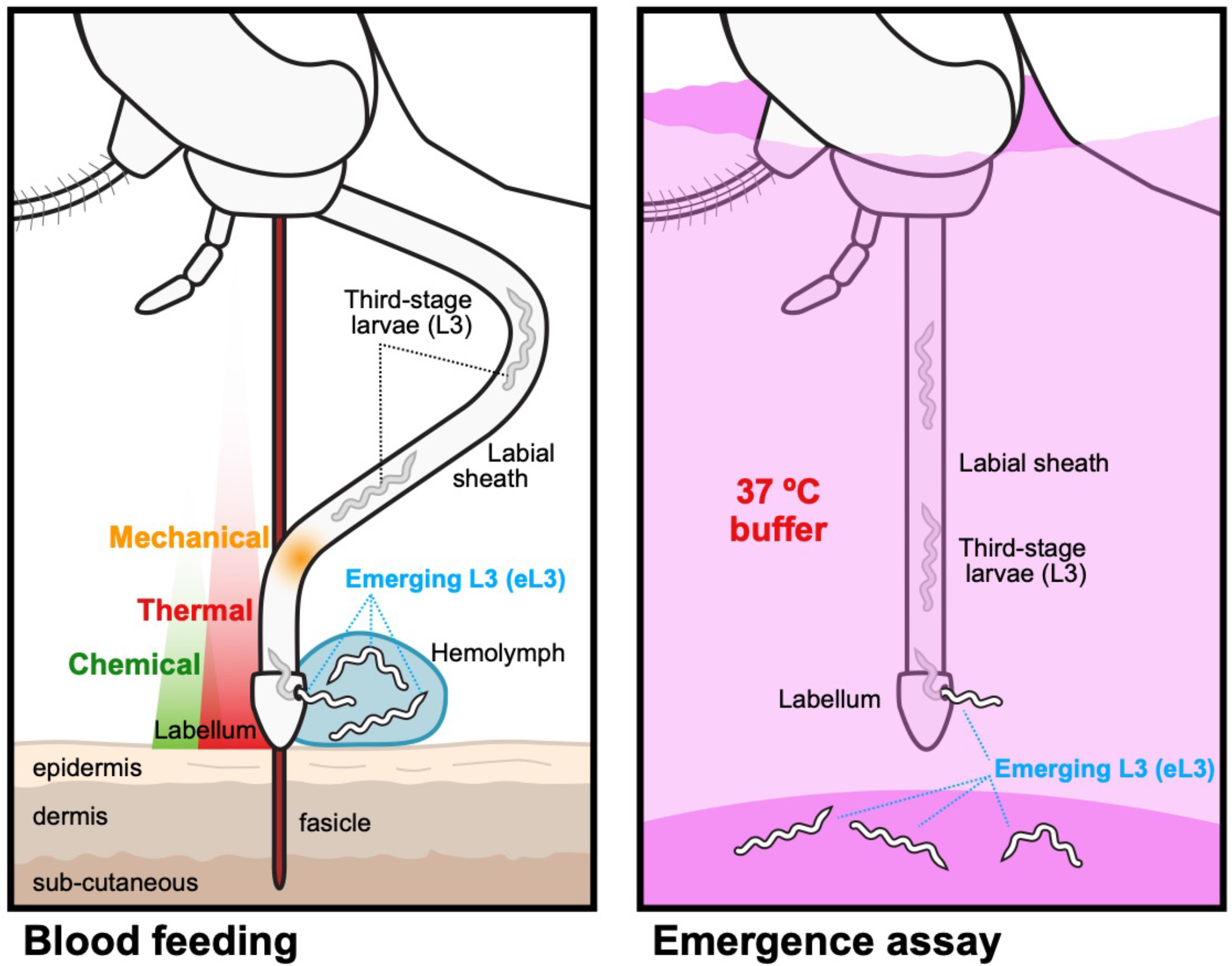

## Background

Arthropods serve as intermediate hosts and vectors of numerous human and animal infective filariae that contribute to a large disease burden worldwide. Indeed mosquito-transmitted lymphatic filariasis, caused by *Wuchereria bancrofti, Brugia malayi* and *Brugia timori*, affects approximately 120 million people in 83 countries [1]. Mosquitoes are also responsible for transmission of animal infective filariae, the most studied being *Dirofilaria immitis*, the agent of canine heartworm disease. Although detailed global numbers are not available, there are an estimated 250,000-500,000 infected dogs in the United States with some areas, such as the Mississippi River basin, reporting infection prevalence as high as 40% [2, 3]. Humans can also be incidentally infected by *D. immitis*, but they do not support the entire life cycle and typically present with only mild clinical signs [3].

Within the mosquito host, ingested microfilariae migrate from the midgut to specific tissues particular to the species. The filarial agents of lymphatic filariasis migrate to the indirect flight muscles in the thorax whereas *D. immitis* migrates into the Malpighian tubules [4]. In these tissues, the parasites develop intracellularly, undergoing successive molts to form third-stage larvae (L3). Some L3 migrate to the proboscis where they are poised for transmission (supplemental movie 2). Infective third-stage larvae emerge most typically from the mosquito labellum but can also emerge from the labial sheath during blood feeding (Fig. 1) [5]. Larvae emerging from the mosquito are deposited on the skin in a drop of hemolymph where they can enter the skin through the bite site [5].

**Fig. 1.**
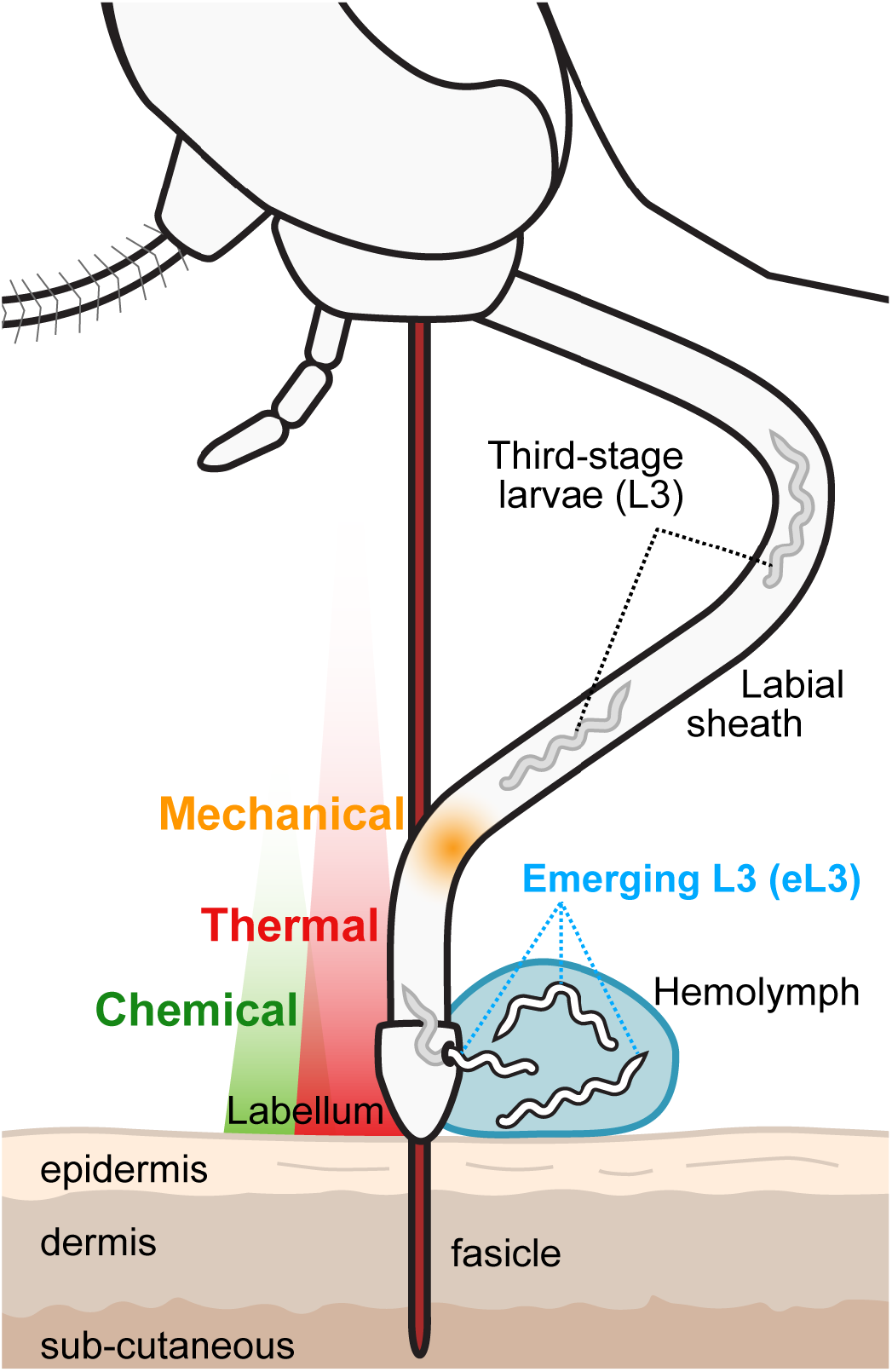
Emergence of infective L3 from tip of the mosquito proboscis. During blood feeding, a subpopulation of L3 in the labial sheath of the proboscis, emerge from the proboscis, alighting on the skin of the host in a drop of the mosquito hemolymph (eL3). Emergence from the proboscis typically occurs at the labellum or distal portion of the labial sheath and can be triggered by sensation of a thermal cue (red gradient). Other cues may also play a role, such as chemical compounds released by the host (green gradient), or by sensation of mechanical forces caused by the deformation of the labial sheath (orange gradient), which slides backward as the fascicle is inserted into the host skin. This figure is adapted from Bancroft B Med J, 1904 [27].

Due to their importance to the disease cycle, much attention is focused on the transmission of infective L3. Different methods have been employed to assay L3 in mosquitoes. A common approach, especially for field isolates, is PCR detection of larvae in the head and other body parts following dissection [6, 7, 8]. Though this method is not established for all filariae, stage-specific PCR has been used for detection of infectious L3 in *W. bancrofti* infected mosquitoes [9]. PCR is sensitive, species-specific, and can be used on pools of mosquitoes in cases where the infection prevalence is low. Another commonly used approach is to physically examine larvae in the labial sheath of the proboscis or in dissected tissues of the mosquito. This approach is particularly useful in a laboratory setting to determine the number of larvae at each developmental stage and to identify their tissue of residence. However, while each of these assays can reveal the number of L3 that have developed, specific assays to enumerate L3 capable of emerging when the mosquito blood feeds have not been previously described.

In addition to assessing the intensity of transmission in field populations, an ability to enumerate and collect L3 larvae capable of emergence would greatly enhance functional assessment. Currently, studies of host immune responses to L3, as well as in vitro and in vivo drug efficacy studies, are typically carried out on L3 collected from infected mosquitoes gently disrupted with a mortar and pestle and then separated using mesh [10, 11, 12]. However, it is unknown whether all L3 harvested by this method are mature and competent to infect the host. Blocking parasite development in the vector is a novel approach being considered for controlling disease transmission and requires a thorough understanding of the molecular interactions between parasites and their vector. The ability to quantify the prevalence of mosquitoes with emerging L3 and determine the number of emerging L3 per mosquito would greatly facilitate this work. Here we present a new assay to quantify emerging infectious L3 that works with different vector and filarial species. We then use this assay to study the development and characteristics of emerging L3 *D. immitis* in mosquitoes. The detailed infection parameters and assay conditions presented here to quantify and produce maximum yields of emerging L3 could potentially facilitate both studies between filariae and their mosquito and vertebrate hosts.

## Methods

### Mosquito strains and culture

*Ae. aegypti* and *Ae. albopictus* strains were provided by the NIH/NIAID Filariasis Research Reagent Resource Center (FR3) and the Malaria Research and Reference Reagent Resource Center (MR4) for distribution by BEI Resources, NIAID, NIH. *Ae. aegypti^S^* (*Ae. aegypti*, Strain Black Eye Liverpool, Eggs, FR3, NR-48921) is a *D. immitis* and *B. malayi* susceptible strain. *Ae. aegypti^R^* (*Ae. aegypti*, Strain LVP-IB12, Eggs, MR4, MRA-735, contributed by David W. Severson) is a *D. immitis* and *B. malayi* refractory strain. Both strains were reared at 27 °C and 80% humidity with a 12-hour photoperiod. *Ae. albopictus^NJ^* (*Ae. albopictus* Strain ATM-NJ95, Eggs, MR4, NR-48979) [13] was reared at 24 °C and 70% humidity with 16:8-hour photoperiod. Mosquitoes were housed in 30 cm^3^ cages (Bugdorm, Taiwan) at a density of approximately 1000 per cage. Larvae were maintained at a density of 1 larva/3 mL. Larvae were fed a suspension of liver powder in water (MP Biomedicals, Solon, OH) and adults were maintained with 10% sucrose in water changed daily. Heparinized sheep blood (Hemostat, Dixon, CA) was provided using an artificial membrane feeder at 37 °C for egg production.

### Mosquito infection with Dirofilaria immitis and Brugia malayi

A full protocol with greater detail is available online (dx.doi.org/10.17504/protocols.io.2xggfjw) [14]. Blood containing *D. immitis* microfilariae was obtained from an experimentally infected dog according to Institutional Animal Care and Use Committee approved protocols and in accordance with the guidelines of the Institutional Animal Care and Use Committee of the University of Pennsylvania (IACUC, protocol 805059). The microfilaremia in this dog is approximately 50,000 mf/mL. Blood containing *B. malayi* microfilariae (*B. malayi* microfilariae in cat blood, live, FR3, NR-48887) was obtained from an experimentally infected cat containing approximately 12,000 mf/mL provided by the NIH/NIAID Filariasis Research Reagent Resource Center for distribution by BEI Resources, NIAID, NIH. In both cases, blood was diluted with the appropriate volume of heparinized sheep blood to a concentration of 4000 mf/mL. The sample was warmed to 37 °C and 3.5 mL was placed in the indentation on the outside of the bottom of a 300 mL plastic baby bottle (Advent, Phillips, Amsterdam, NL) and covered with Parafilm. The inside of the bottle was filled with water at 37 °C. This assembly was placed with the membrane side down on the top mesh of mosquito cages and the insects are allowed to feed for 15 minutes (Fig. 2A). Immediately following the feed, blood fed mosquitoes were separated using CO2 anesthesia and housed under standard insectary conditions described above.

**Fig. 2.**
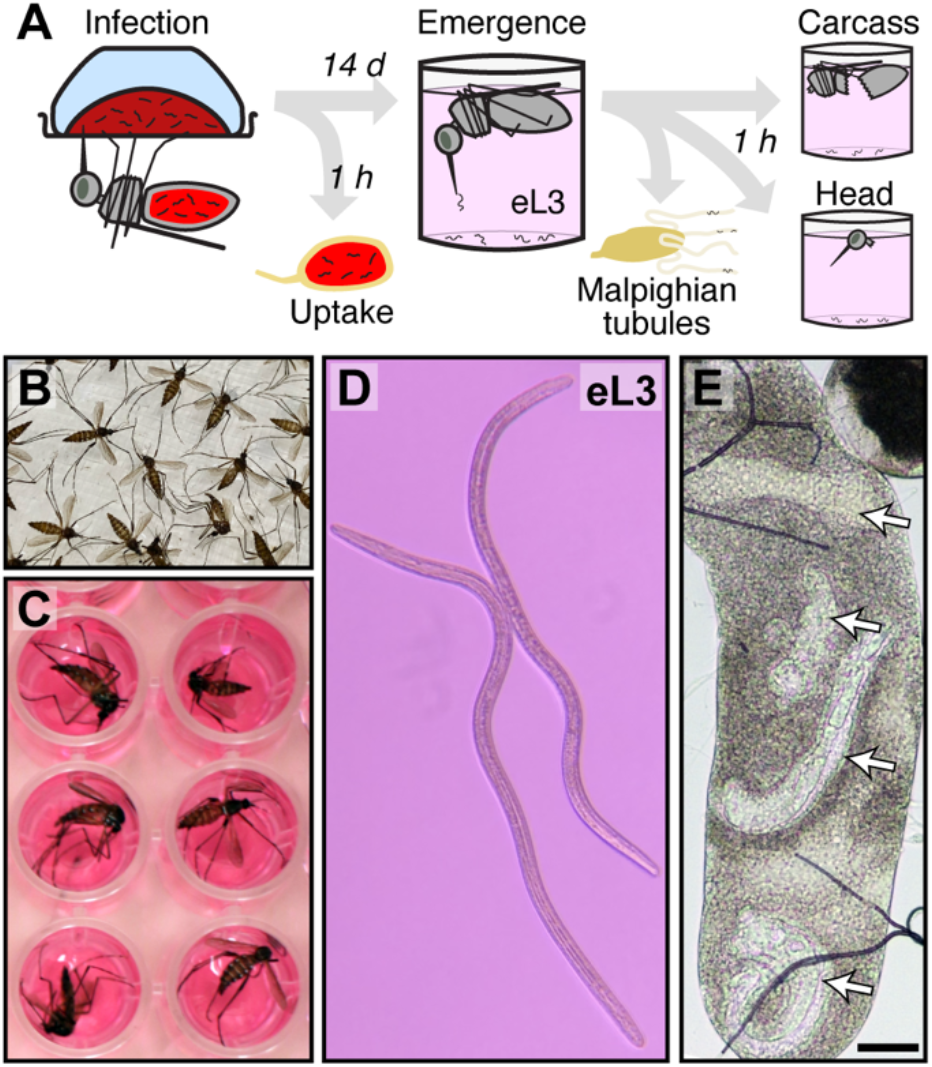
Assay for eL3. (**A**) Infection and assay protocol followed in this study. Mosquitoes are fed on blood containing *D. immitis* microfilariae. Uptake is measured in a small group, and the remaining mosquitoes are maintained until an emergence assay is performed. Immediately following the emergence assay with whole mosquitoes, the Malpighian tubules are dissected and analyzed. The dissected head and carcass are individually placed into an emergence assay to assay L3 that failed to emerge from intact mosquitoes. Figure S1 shows images of dissected mosquitoes and Malpighian tubules. (**B**) Mosquitoes are rinsed with 70% ethanol to wet, rinsed with water, and (**C**) placed individually into wells of a 96-well plate. (**D**) Emerging third-stage larvae (eL3) from intact mosquitoes or L3 from dissected heads and carcasses at the bottom each well are scored by microscopy. Supplemental movie 1 shows typical movement of eL3. (**E**) Larvae (white arrows) in live Malpighian tubules are scored by microscopy. Scale bar is 50 μm.

### Microfilarial uptake

Within an hour of blood feeding, fed mosquitoes were anesthetized with carbon dioxide at room temperature. The entire undamaged midgut was dissected in deionized water in a depression slide and immediately transferred to a standard microscope slide containing a 50 μL drop of deionized water. Damaged midguts leaking blood during dissection were not used. The epithelium of the gut was separated from the blood bolus and moved through the water to remove any residual blood before it was discarded. The blood was roughly dispersed using forceps and then pipetted up-and-down with a 20 μL pipette. Microfilaria in the entire drop were counted immediately without a coverslip.

### Emergence assay for eL3 and other larval stages

A full protocol with greater detail is available online (dx.doi.org/10.17504/protocols.io.xbyfipw) [15], which is a modified from a previously published method [16]. Briefly, eL3 were assayed by placing live mosquitoes in 70% ethanol in water for 1 min. Afterward the mosquitoes were rinsed twice in deionized water and then placed individually into wells of a 96-well plate containing 200 μL Dulbecco’s Modified Eagle’s Medium (DMEM) with L-glutamine, high glucose and sodium pyruvate (Corning Mediatech, Manassas, VA). At this point, the mosquitoes are still alive, and can be observed moving in the buffer, but they cannot stand on the surface due to the wetting procedure. The plate was placed in a 37 °C incubator with 5% CO2 for 60 minutes and the number of eL3 was determined without removing the mosquitoes using an inverted microscope with a 4x objective. In some experiments, after the eL3 emergence assay, mosquitoes were removed from the plate individually for assaying other larvae. First, the Malpighian tubules were dissected by removing the posterior two abdominal segments. The set of five Malpighian tubules was removed from the midgut and transferred to a microscope slide containing 15 μL PBS and covered with a 22 mm^2^ coverslip. The larvae were counted and although different stages were present, we did not categorize them (Fig. S2). The head was separated from the body. The dissected head and carcass fragment were placed separately into fresh wells of a 96-well plate prepared and incubated as described above. L3 larvae were allowed to migrate out of the dissected tissue for at least 60 minutes and scored as described above. Only a portion of the mosquitoes used for the eL3 emergence assay were processed in this manner.

### Emergence assay time course and microfilaria concentration series

Groups of 300-500 mosquitoes were fed on blood containing *D. immitis* microfilariae at different concentrations and housed as described above. Mosquito mortality was monitored daily as assessed by the fraction of the population that died each day. This was necessary since groups of 50 mosquitoes were removed for eL3 assays as described above. Since the emergence prevalence was very low on day 12, emergence assays were performed with only 25 mosquitoes per replicate saving more of the population for later days where there is a higher eL3 prevalence. For concentration series experiments, a blood dilution containing 32,000 mf/mL was created and then subjected to two-fold serial dilution to create feeding doses of 16,000 mf/mL, 8000 mf/mL, and 4000 mf/mL. For each concentration, five mosquitoes of each strain were used to measure uptake as described above. For these experiments, mortality was monitored daily until day 17 when an assay for eL3 was performed on the remaining mosquitoes. Kaplan-Meier survival analysis was performed between adjacent doses within each mosquito strain and between the same doses across the mosquito strains.

## Results

### Novel assay for filarial nematode L3 emerging from individual mosquitoes

In our assay, outlined in Fig. 2A, we infect mosquitoes using an artificial membrane feeder, and determine the number of microfilariae ingested immediately afterwards [14]. The microfilarial uptake represents the theoretical maximum number of parasites in the assay. To assay the L3 capable of emerging from the mosquitoes, we simulate the thermal cue that would be experienced by a mosquito landing on a mammalian vertebrate host by placing mosquitoes in buffer and warming to 37 °C [15]. Mosquitoes are placed individually into wells of a 96-well plate after the wetting procedure (Fig. 2B-C), and care is taken to minimize damage to the mosquito during transfer. Upon warming, competent L3 emerge from the mosquito and sink to the bottom of the well, where they can be counted (Fig. 2D and supplemental movie 1). Emerging L3 larvae (hereafter referred to as eL3) collected by this method are capable of molting to the L4 stage in vitro [17] and are infectious to dogs [16]. After the emergence assay, the mosquitoes are still viable and can move, but cannot escape the buffer due to the wetting procedure. Notably, following the emergence assay, further analyses can be carried out to determine the number and location of larvae remaining within the mosquito. In our study, both dissected Malpighian tubules (Fig. 2E), and the head and remaining carcass were assayed separately as described below (Fig. S1).

### Microfilariae robustly migrate into the Malpighian tubules but only a fraction develop to eL3

To determine the efficiency of eL3 development we infected *D. immitis* susceptible (*Ae. aegypti^S^*) or refractory (*Ae. aegypti^R^*) strains and performed an emergence assay 14 days post-infection. Consistent with their resistant phenotype, we found that *Ae. aegypti^R^* released no eL3, despite having a median uptake of 12 microfilariae per mosquito (Fig. 3A). In the susceptible strain (*Ae. aegypti^S^*), 81 of the 252 mosquitoes assayed (32%) had at least one eL3 when given a median uptake of 15 microfilariae per mosquito (Fig. 3B). A portion of the susceptible mosquitoes used for the emergence assay were dissected and a substantial number had L3 remaining in the dissected heads (26 of 102; 25%) and carcasses (48 of 102; 47%) (Fig. 3B). The majority of larvae assayed at day 14 were present in the Malpighian tubules and nearly all of the dissected mosquitoes (93 of 96; 97%) had at least 1 larva present in the Malpighian tubules (Fig. 3B). In a few cases, Malpighian tubules were damaged during the dissection and were excluded from this and subsequent analyses. Pooling all larvae present across all tissues revealed that the median total number of larvae per mosquito was 8. Therefore, at day 14 post infection, 53% of the 15 ingested microfilariae could be accounted for as larvae in the mosquito (Fig. 3B). In the dissected Malpighian tubules, we found larvae in various stages of development, but we did not determine their stage (Fig. S2). We note however, that larvae found in the Malpighian tubules, even the most stunted sausage forms, are viable, as they were capable of movement. Previous studies suggest that not all L3 emerge when infected mosquitoes blood feed on an animal [5]. Since this previous observation was not quantitative, we analyzed the data from the emergence assay (Fig. 3B) to compare the number of eL3 to the total L3 (eL3 plus L3 in dissected heads, and dissected carcasses). We did not include larvae found in the Malpighian tubules since we could not unambiguously characterize them as L3. We found that the number of eL3 is significantly lower than the total L3 (Fig. 3C). Therefore, only a fraction of the larvae that develop to L3 emerge during our emergence assay, consistent with previous observations [5].

**Fig. 3.**
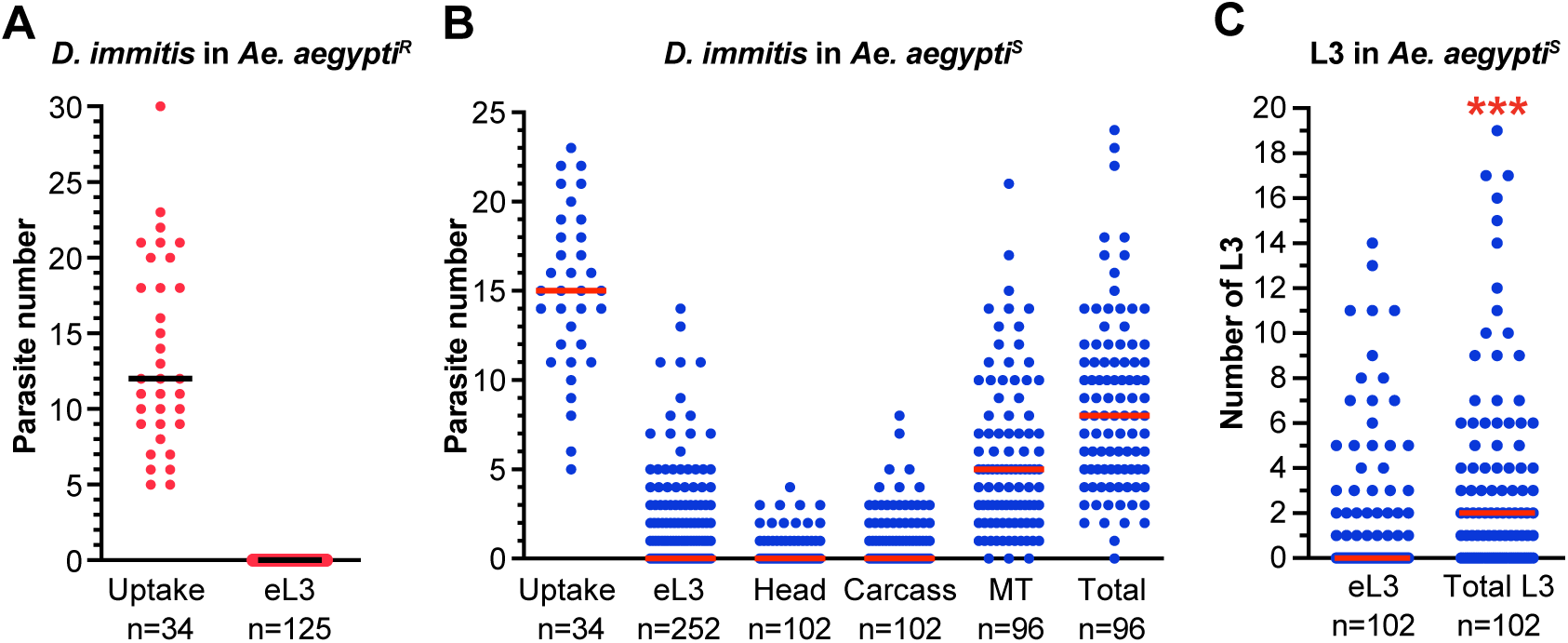
A fraction of ingested microfilariae develops into eL3 in *Ae. aegypti^S^*. (**A**) Dots indicate the number of microfilariae present in the midgut of individual *Ae. aegypti^R^* immediately after feeding on infected blood (Uptake) and the number of eL3 assayed 14 days post infection. The black line indicates the median. (**B**) Dots indicate the number of microfilariae present in the midgut of individual *Ae. aegypti^S^* immediately after feeding on infected blood (Uptake), the number of eL3 assayed 14 days post infection, the number of L3 emerging from the dissected heads (Head) or carcass (Carcass), and the number present in the Malpighian tubules (MT). The Total is sum of all parasites of any stage found in any tissue or assay on day 14. The red line indicates the median. (**C**) Dots are the number of eL3 from individual *Ae. aegypti^S^* and all third-stage larvae assayed (Total; sum of eL3, Head, and Carcass) (data taken from panel B). The red line is the median number. The asterisks indicate a Mann-Whitney P-value < 0.001. These data are the sum of three independent biological replicates. The number of mosquitoes assayed is shown below each column.

### Once eL3 have developed, their intensity and prevalence are constant

To determine the optimal day to perform the emergence assay, we wanted to identify a time point at which eL3 recovery was maximized, but mosquito mortality was still minimal. To do this, we infected large populations of mosquitoes and then removed groups of approximately 50 mosquitoes on consecutive days for an emergence assay. Four independent experiments were pooled since it was not possible to obtain all time points from a single infection. We assessed the variation in eL3 number over the 10-day period from day 12-21 post infection. The main emergence phase started at day 13, as we observed only a single eL3 in 48 mosquitoes at day 12 post-infection. Once L3 begin to emerge from mosquitoes, both the number that emerge per mosquito as well as the prevalence of mosquitoes with at least one emerging L3 are relatively constant from days 13-21 post infection (Fig. 4). These data suggest that larvae develop synchronously and that once they reach the L3 stage, their numbers remain relatively stable over time. Beyond day 13, mosquito mortality continues and eL3 prevalence or intensity is not increased, Therefore, to maintain population sizes and maximize eL3 yield we typically assay mosquitoes 14-17 days post infection.

**Fig. 4.**
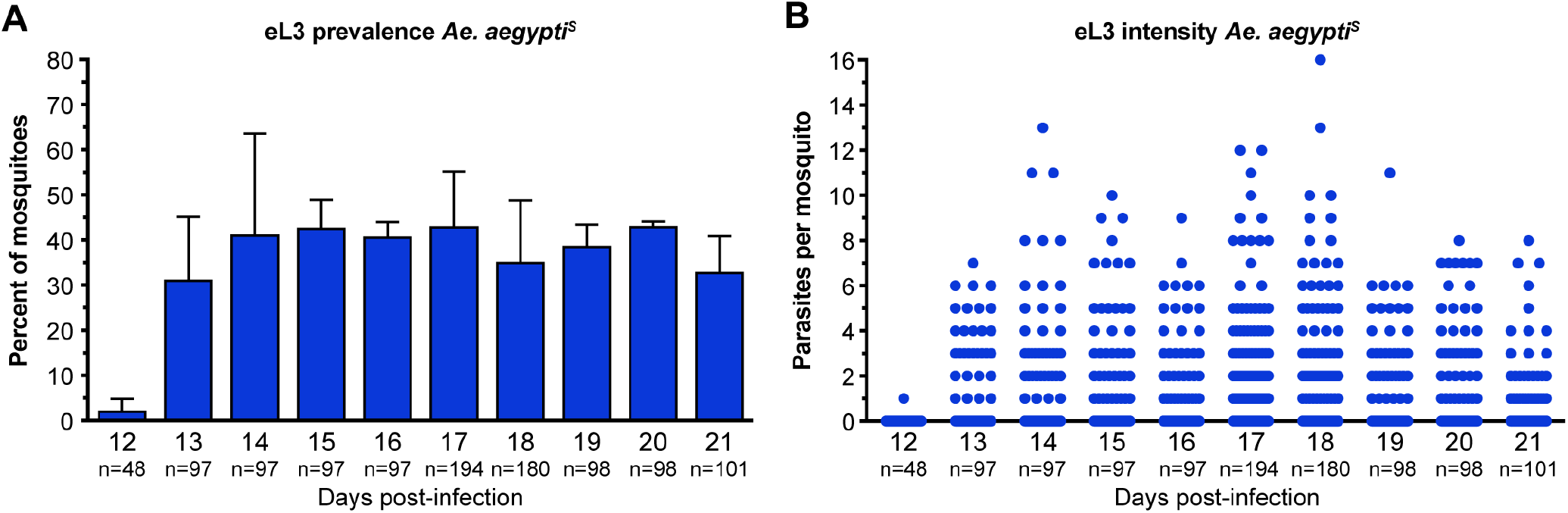
The number of eL3 *D. immitis* from *Ae. aegypti^S^* are constant 13-21 post infection. (**A**) Graph of the average prevalence in *Ae. aegypti^S^* of *D. immitis* eL3 assayed 12-21 days post-infection. Error bars indicate the standard deviation. (**B**) Dots are the number of eL3 *D. immitis* from individual mosquitoes assayed 12-21 days post-infection. We performed this experiment four times. Two replicates were assayed on days 12-18. The other two replicates were assayed on days 14-21. Each replicate was performed with approximately 50 mosquitoes. The replicates on day 12 were performed with fewer mosquitoes since our preliminary data suggested that emergence was negligible on this day.

### Increasing the number of D. immitis microfilariae ingested does not increase eL3 prevalence

To optimize the number of eL3 recovered from the mosquito, we next determined if increasing the number of microfilariae in the blood meal improved the recovery of eL3. For these experiments we used the standard concentration of 4000 mf/mL and included three additional concentrations of 8000, 16,000, and 32,000 mf/mL. We found that increased concentrations of microfilaria in the membrane feeder had a consequent impact on the number present in the mosquito midgut immediately after blood feeding, with no significant difference between *Ae. aegypti^S^* and *Ae. aegypti^R^* (Fig. 5A). At each successive concentration, microfilariae present in the blood meal increased approximately two-fold. We monitored mosquito survival for 17 days, at which time an emergence assay was performed. We chose day 17 to allow more time to monitor mosquito survival. The prevalence of eL3 in *Ae. aegypti^S^* was not significantly different, regardless of the concentration of microfilariae fed (Fig. 5B). On a population level, the number of eL3 produced per mosquito was not increased by feeding concentrations of microfilariae greater than 4000 mf/mL (Fig. 5C). Increasing the concentration of *D. immitis* microfilaria used to infect *Ae. aegypti^S^* increased mosquito mortality in a dose-dependent manner (Fig. 5D). Each increase in microfilaria concentration significantly increased mortality from the previous one, including the lowest concentration, which elevated mortality compared to those fed on uninfected blood. A similar dose-response was observed in the refractory strain; however, only the uninfected blood to 4000 mf/mL and 16,000 to 32,000 mf/mL comparisons were significantly different (Fig. 5E). When we compared the survival of *Ae. aegypti^S^* and *Ae. aegypti^R^* at the different concentrations of microfilariae, there was no difference in survival at the lowest concentration (Fig. S3). There was a modest difference at 8000 mf/mL, with *Ae. aegypti^S^* showing greater mortality, and this difference further increased at the two highest concentrations.

**Fig. 5.**
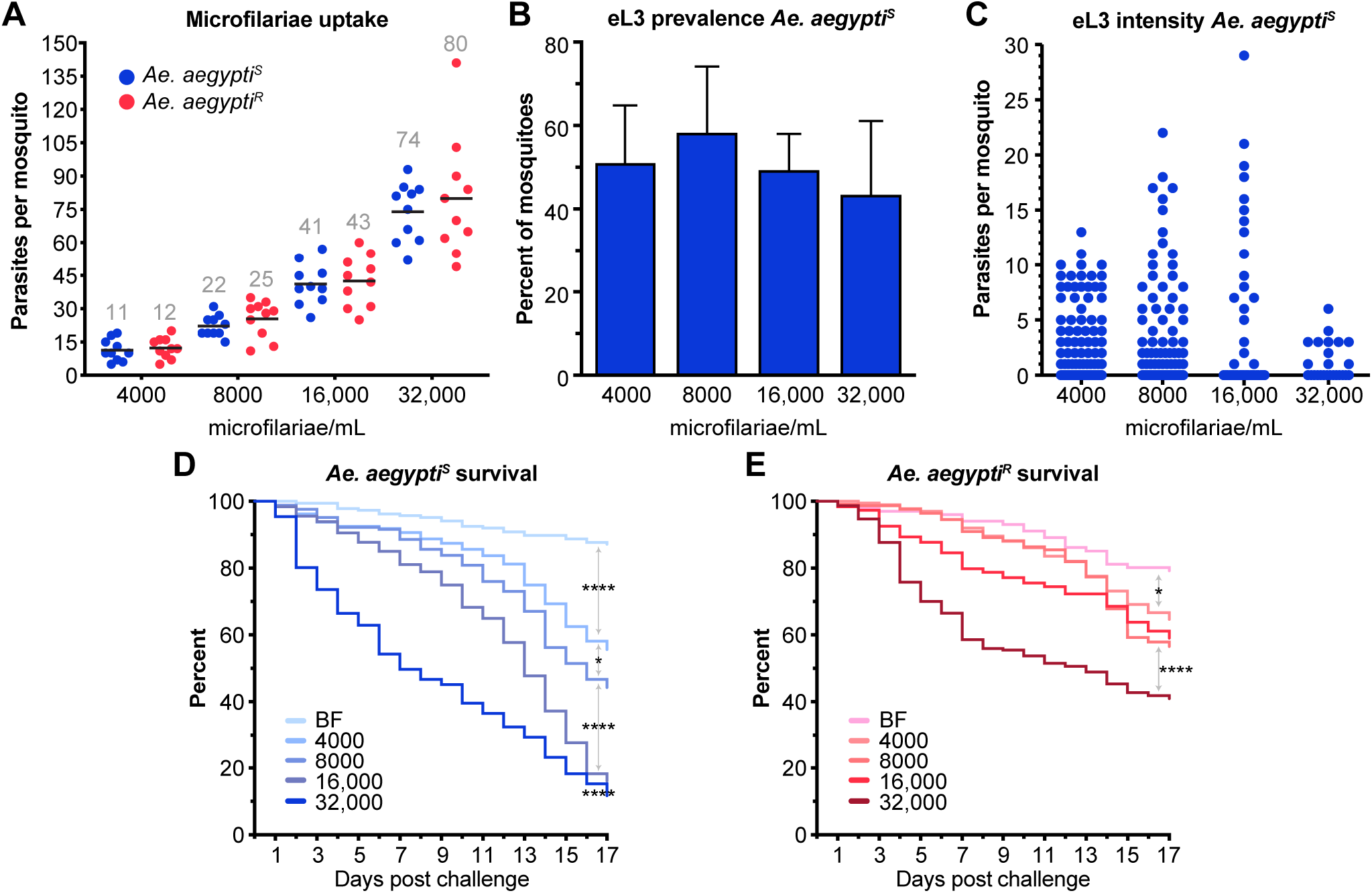
Increasing the dose of microfilariae does not increase numbers of eL3 but increases mosquito mortality. (**A**) Dots indicate the number of *D. immitis* microfilariae present in midguts of *Ae. aegypti^S^* (blue) and *Ae. aegypti^R^* (red) immediately following blood feeding on the indicated doses of microfilariae. Data in each column are normally distributed and black lines and numbers indicate the mean. No significant differences were found when *Ae. aegypti^S^* and *Ae. aegypti^R^* were compared at the different doses using an unpaired t test. Data are from two independent biological replicates. (**B**) Graph of the average prevalence of *D. immitis* eL3 in mosquitoes feeding on blood with increasing concentrations of microfilariae, assayed 17 days post-infection. Error bars indicate the standard deviation. There was no significant difference comparing the columns using and ANOVA. (**C**) Dots are the number of *D. immitis* eL3 from individual mosquitoes assayed 17 days post-infection. No significant differences were found when we compared all groups to each other or comparing 4000 mf/mL to the other groups using a Kruskal-Wallis test with Dunn’s correction for multiple comparisons. Data for panels B and C are pooled from four separate biological replicates. (**D**) Kaplan Meier survival plot for *Ae. aegypti^S^* or (**E**) *Ae. aegypti^R^* fed with uninfected blood (BF) or blood containing different concentrations of microfilariae. Pairs of adjacent treatment groups were analyzed by Kaplan Meier, and relationships with significant differences indicated with asterisks in Figure S3 and in Table S1.

### Stored blood containing microfilariae retains its ability to infect mosquitoes and produce eL3

All *D. immitis* experiments described above used freshly isolated microfilaremic blood. However, there are times when it is not possible to use this material immediately, such as when blood is shipped from another location. We wanted to compare the number of eL3 produced when mosquitoes were fed infected blood stored at 4 °C for one or two days. For these experiments, we fed mosquitoes with freshly isolated infected blood and on the next two consecutive days fed new cohorts with the same sample that had been stored at 4 °C. We performed an emergence essay at day 17 of each infection. There was no significant difference in the prevalence of eL3 between mosquitoes fed fresh microfilaremic blood and ones fed on an aliquot of the same blood sample stored for one day at 4 °C (Fig. 6A). However, there were significant decreases in eL3 prevalence in mosquitoes fed on microfilaremic blood stored for two days at 4° C compared to mosquitoes fed on blood stored at 4° C for one day and to mosquitoes fed on fresh blood. Similar trends were observed in infection intensity (Fig. 6B). Our data show that although microfilaremic blood stored at 4 °C for 2 days can still infect mosquitoes, there is a trend towards lower numbers of eL3 following blood storage, suggesting that refrigerated samples should be used as soon as possible for maximum infection potency.

**Fig. 6.**
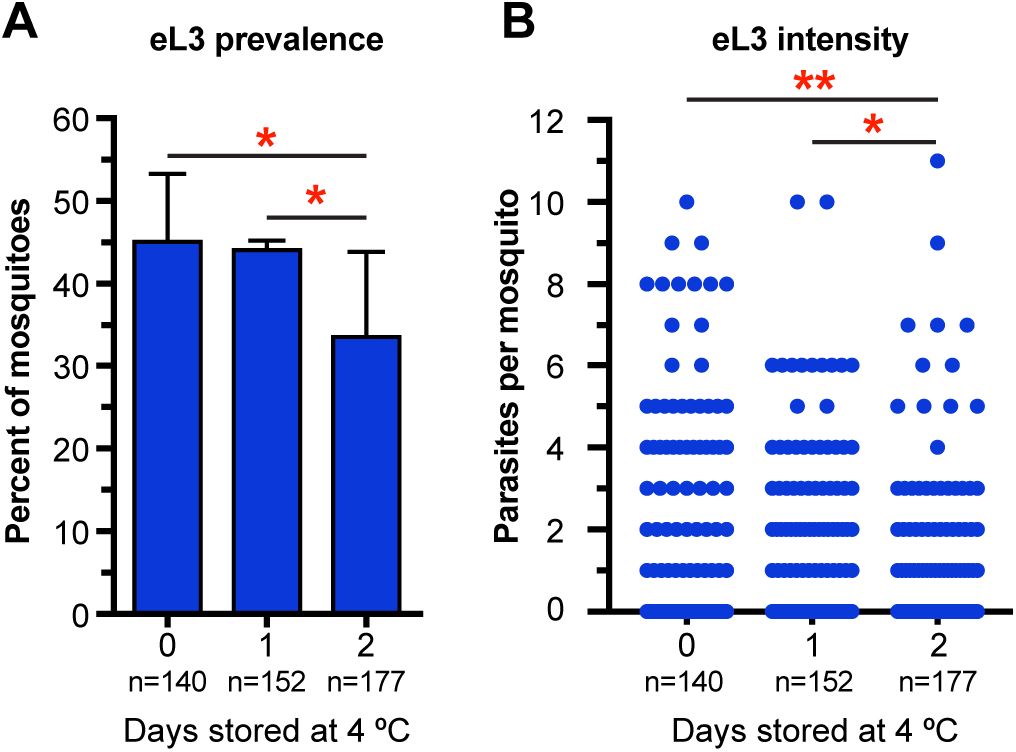
Infected blood stored at 4 °C produces eL3. Infected blood was used fresh (0) or after storage for 1 or 2 days at 4°C. (**A**) Graph of the average prevalence of *D. immitis* eL3 17 days postinfection in *Ae. aegypti^S^.* The error bar indicates the standard deviation. The number of mosquitoes (n) analyzed indicated for each sample. The asterisks indicate P<0.05 using a Fisher’s exact test. (**B**) Blue dots are the number of *D. immitis* eL3 emerging from individual mosquitoes assayed 17 days postinfection. Significant differences in intensity using Kruskal-Wallis test with Dunn’s correction for multiple comparisons are indicated with one and two asterisks for P<0.05 and <0.01, respectively. Data pooled from two independent experiments.

### Emergence assay works in natural vector of Dirofilaria immitis

Since *Ae. aegypti* is not considered an important natural vector of *D. immitis*, we wanted to determine whether eL3 could be obtained from a mosquito species associated with *D. immitis* transmission. For these experiments we used a strain of *Ae. albopictus* isolated from Keyport, New Jersey (*Ae. albopictus^NJ^*). We found that eL3 were present at day 17, but their prevalence and number were lower than those present in *Ae. aegypti^S^* (Fig. 7A-B) when the mosquitoes were reared and housed in our standard insectary conditions (27 °C with 80% relative humidity and a 12:12 h photoperiod). When we tested *Ae. albopictus^NJ^* reared and infected at its ideal temperature and humidity (24 °C, 70% relative humidity, and a 16:8 h photoperiod), the prevalence and number of eL3 were increased, although they were still lower than typical for *Ae. aegypti^S^* (Fig. 7C-D). This is likely because the strain of *Ae. albopictus^NJ^* used did not feed to repletion from an artificial membrane feeder under any conditions that we tested. Even though we are not able to make quantitative comparisons to *Ae. aegypti^S^*, our data show that the emergence assay works with different mosquito species.

**Fig. 7.**
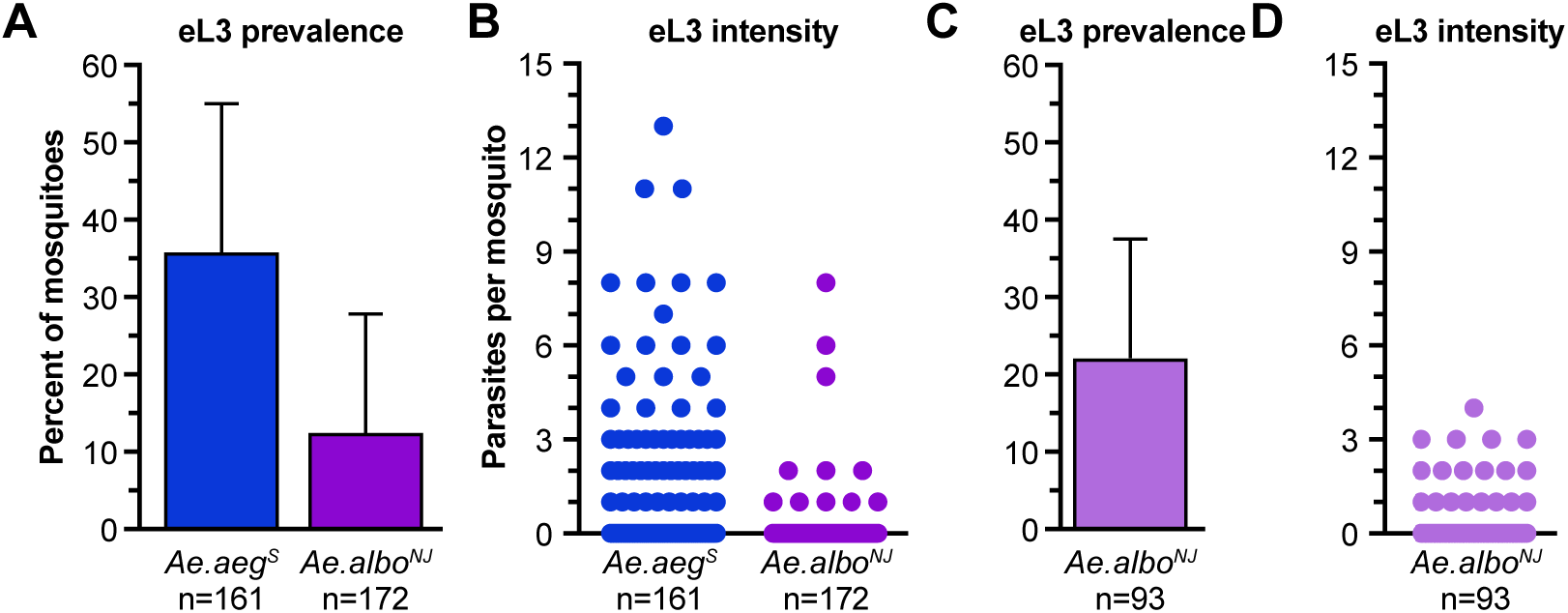
*Aedes albopictus* supports eL3 development. (**A**) Graph of the average prevalence of *D. immitis* eL3 17 days post-infection in *Ae. aegypti^S^* (blue, *Ae. aeg^S^*) or *Ae. albopictus^NJ^* (purple, *Ae. albo^NJ^*). Error bars indicate the standard deviation. The number of mosquitoes (n) analyzed indicated for each sample. (**B**) Blue and purple dots are the number of eL3 *D. immitis* larvae emerging from individual *Ae. aegypti^S^* (*Ae. aeg^S^*) or *Ae. albopictus^NJ^* (*Ae. albo^NJ^*) mosquitoes assayed 17 days postinfection, respectively.

### Individual mosquito emergence assay for transmission stage Brugia malayi

To determine whether the emergence assay will be informative for enumerating eL3 for other filarial nematodes, we infected mosquitoes with *B. malayi*. These parasites have a shorter developmental period in the mosquito [18], so we performed our emergence assay on days 12-14 post infection in both *Ae. aegypti^S^* and *Ae. aegypti^R^* mosquitoes. Despite taking up a similar number of microfilariae in the blood meal (Fig. 8A), no eL3 were produced by *Ae. aegypti^R^*, showing that this strain is refractory to both *B. malayi* and *D. immitis* (Fig. 8B). In contrast, a significant proportion of *Ae. aegypti^S^* mosquitoes develop eL3, showing that this strain is susceptible to both *B. malayi* and *D. immitis* (Fig. 8B). There was no significant difference in either the prevalence of mosquitoes producing eL3 or in the number of eL3 produced per mosquito between the days we assayed (Fig. 8B-C). These data indicate that despite their different lifecycle in the mosquito, eL3 of human filariae can be measured using our emergence assay.

**Fig. 8.**
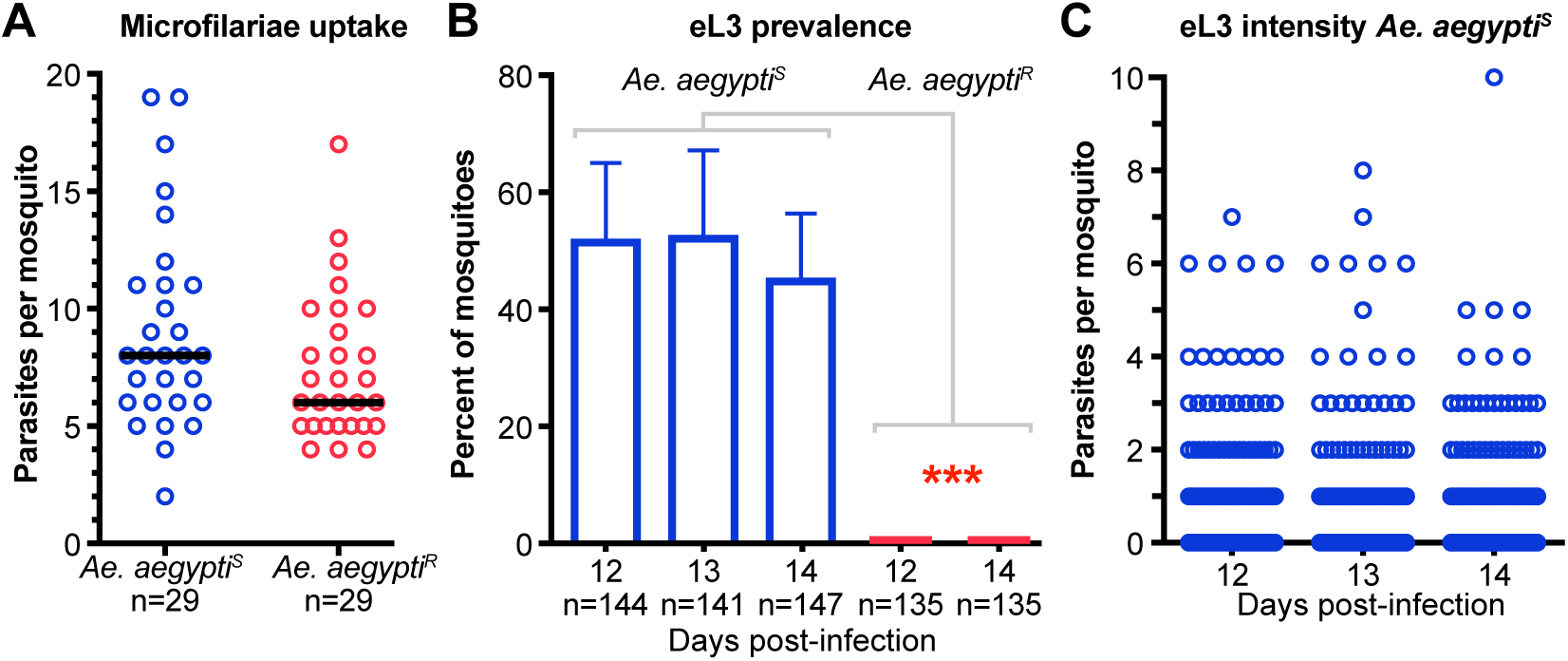
Emergence assay can be used to quantify *B. malayi* eL3. (**A**) Dots indicate the number of *B. malayi* microfilariae present in midguts of *Ae. aegypti^S^* (blue) and *Ae. aegypti^R^* (red) immediately following blood feeding. Data in each column are not normally distributed and black line is the median. The number of mosquitoes (n) analyzed indicated for each sample. No significant difference was using a Mann Whitney test (P=0.09). (**B**) Graph of the average prevalence of *B. malayi* eL3 assayed 12-14 days post-infection in *Ae. aegypti^S^* (blue bars) and at days 12 and 14 in *Ae. aegypti^R^* (red bars). Error bars indicate the standard deviation. The number of mosquitoes (n) analyzed indicated for each sample. No emerging parasites were observed for *Ae. aegypti^R^*. There were no significant differences between the days assayed for *Ae. aegypti^S^*, and all *Ae. aegypti^S^* days assayed were significantly different from both *Ae. aegypti^R^* on the two days assayed using an ANOVA with Tukey’s multiple comparisons test as indicated by asterisks (P<0.001). (**C**) Blue dots are the number of *B. malayi* eL3 from individual mosquitoes assayed 12-14 days post-infection. No significant differences were found when we compared all groups to each other or comparing day 12 to the other groups using a Kruskal-Wallis test with Dunn’s correction for multiple comparisons. Data were pooled from three independent experiments.

## Discussion

Development of an assay for eL3 may be valuable for studying mosquito-filaria interactions since it provides a clearer indication of vector competence and vector transmission intensity. As our data and a previously reported observation [5] indicate that not all L3 residing in the mosquito emerge when stimulated either by warming the mosquito or by direct contact with mammalian host, even if already residing in the head, our method provides a more accurate indication of a particular vector species’ potential for pathogen transmission. For example, we have found that not all strains of *Ae. aegypti* or *Ae. albopictus* support eL3 development, even though microfilariae invade the Malpighian tubules. In other studies, we used this assay to show that activation of the Toll immune signaling pathway restricts development of *D. immitis* and *B. malayi* eL3 [19, 20]. In addition, this assay could also permit assessment of manipulations of the mosquito host or larvae that result in an inability of L3 to emerge in response to thermal stimuli or due to motor defects that could be missed by current methods. Finally, results from our assays suggests that quantitating all L3 by dissection will likely result in overestimating the number of larvae that are capable of emerging from the mosquito. This is particularly true in the case where the number of L3 is derived by mosquito homogenization, as L3 from the body cavity and those present in the Malpighian tubules are likely included.

Results of our experiments to optimize recovery of eL3 by increasing the dose of *D. immitis* microfilariae given to mosquitoes reveal that microfilarial uptake and eL3 emergence are uncoupled, as we did not observe significant increases in the prevalence of mosquitoes with eL3 or the number of eL3 per mosquito at the doses tested. While it is likely that microfilarial uptake and eL3 emergence are coupled at lower concentrations of microfilariae, we did not test this since our main goal was to establish conditions where eL3 recovery is maximized to support future study of transmission-blocking strategies. Varying the timing of the assay revealed that after day 13 there is no significant difference in the prevalence or number of eL3. Taken together, our data suggest that the Malpighian tubules have a limited capacity to support *D. immitis* development and, once the limit is reached, no additional eL3 are produced. However, we observed a range of developmental phenotypes in the Malpighian tubules of mosquitoes 14 days after *D. immitis* infection, raising the possibility that larval damage to the tubules prevents further development, as all larvae are viable at this time point, even those displaying delayed development phenotypes. While, a substantial number of midgut epithelial cells are invaded during infection by ookinetes of *Plasmodium*, the causative agent of malaria, damaged cells are eliminated and replaced through tissue regeneration [21, 22]. As larval development is blocked and mosquito fitness is negatively impacted over time, it suggests that damaged cells of the Malpighian tubules persist and are not replaced. In addition, the continuing metabolism of the increased number of larvae or an accumulation of larval waste products may also contribute to the increased mosquito mortality we observed in *Ae. aegypti^S^* at later timepoints.

By quantifying microfilarial uptake in individual mosquitoes and comparing it to all parasites that emerge from either the whole mosquito, the head, or the carcass or that are present in the Malpighian tubules on day 14, we found that approximately 60% of the ingested microfilariae are accounted for at day 14. Although we are accounting for a majority of the parasites ingested by the mosquitoes, it is nevertheless interesting to speculate about the fate of those that are not assayed. Some L3 remain in the proboscis of dissected heads, even after the primary emergence assay and the subsequent emergence assay after dissection. However, we did not systematically attempt to quantify these, and it remains possible that L3 may also be retained in the dissected carcasses. In addition, because some microfilariae may be damaged by the mosquito immune system or fail to follow sensory cues, they may fail to invade the Malpighian tubules and may be voided into the hindgut and then the feces [4, 23, 24]. Likewise, the activity of microfilariae and larvae in the Malpighian tubule cells may damage these cells, sometimes to the point where microfilariae are released into the tubule lumen, where they also could be eliminated as waste. Alternatively, if the basal side of the Malpighian tubule cell is compromised, parasites could be released into hemolymph within the mosquito body cavity, where they could be eliminated or sequestered. As the possibilities outlined above are not mutually exclusive, we believe they could collectively account for the differential between microfilarial uptake and total parasites ultimately detected.

We used our emergence assay to analyze the effects of storing microfilaremic blood at 4°C and showed that it is possible to produce eL3 after storage for at least two days, but likely even longer. However, while delays in initiating mosquito infection do not preclude experimentation, the strongest prevalence and intensity of eL3 production was observed in mosquitoes infected by ingestion of fresh microfilaremic blood. Given the loss of eL3 production with blood storage time, it is not recommended to make comparisons within an experiment using a blood sample stored for different lengths of time. However, internal comparisons are reasonable, as is the goal of using the stored blood to produce eL3.

Because L3 obtained in our assay have responded to physiological conditions, L3 produced via an emergence assay may provide an easily accessible source of larvae than L3 obtained from gently crushed and sieved mosquitoes [25]. While we developed this emergence assay for use on single mosquitoes, it is also suitable for use on populations en masse and eL3 obtained this way are known to be infective [16]. As such, eL3 may be quite valuable for future identification and testing of novel heartworm preventatives, which requires infectious *D. immitis* L3, as well as for studies of the host immune response to infection, which often require infectious L3. Similarly, in vitro L3 to L4 molting assays, such as the newly developed genetic transformation protocol for *B*. malayi [26], may benefit from L3 produced via our method.

## Conclusions

We have established a novel assay to determine the number of infectious L3 filarial larvae capable of emerging from individual mosquitoes, which we refer to as eL3. We have shown this assay works with both different mosquito and filarial nematode species. As such, our method is likely suitable for assessment eL3 of other mosquito transmitted filariae, such as *D. repens*, a canine and human skin dwelling relative of *D. immitis*, and *Wuchereria bancrofti*, which is responsible for approximately 90% of human lymphatic filariasis and can be transmitted by several genera of mosquitoes (*Culex, Aedes*, and *Anopheles*). It remains to be determined if our assay will work for filariae transmitted by other arthropod vectors such as biting flies, mites, fleas, and ticks. In summary, the method we describe here will greatly facilitate current efforts to block transmission of arthropod transmitted filariae by enabling quantitative analysis in the vector as well as providing a more reliable way to harvest infectious larvae to evaluate new preventatives, treatments, or vaccines.

## Abbreviations

(BE, susceptible, *Ae. aegypti^S^*): *Aedes aegypti* Blackeye Liverpool strain
(LVP, refractory, *Ae. aegypti^R^*): *Aedes aegypti* Liverpool strain
(mf): microfilariae
(MT): Malpighian tubules
(L3): third-stage larvae
(eL3): emerging third-stage larvae

## Declarations

### Ethics approval and consent to participate

Blood containing *D. immitis* microfilariae was obtained from an experimentally infected dog according to Institutional Animal Care and Use Committee approved protocols and in accordance with the guidelines of the Institutional Animal Care and Use Committee of the University of Pennsylvania (IACUC, protocol 805059).

### Consent for publication

Not applicable.

### Availability of data and material

All data generated or analyzed during this study are included in this published article and its supplementary information files.

### Author’s Contributions

MP was responsible for the study conception. ARM, EBE, GTO, FMO, and MP were responsible for acquisition of data. ARM, EBE, GTO, FMO, TJN, JBL, and MP were responsible for analysis and interpretation of data. MP and ARM were responsible for drafting and critical revision of the manuscript. EBE, GTO, FMO, TJN, and JBL were responsible for critical revision of the manuscript.

## Acknowledgements

We thank Leslie King who helped with critical reading of the manuscript and the Filariasis Research Reagent Resource center for providing *Brugia malayi* and *Aedes aegypti* Black Eye Liverpool strain.

---

**Table S1.** Pairwise comparisons of mosquito survival following blood feeding on different doses of *D. immitis* microfilariae.

| ***Ae. aegypti*<sup>S</sup>** | **P-value** |
| --- | --- |
| 4k vs Blood feeding | <0.0001 |
| 8k vs 4k | 0.0337 |
| 16k vs 8k | <0.0001 |
| 32k vs 16k | <0.0001 |
| ***Ae. aegypti*<sup>R</sup>** | **P-value** |
| 4k vs Blood feeding | 0.0142 |
| 8k vs 4k | 0.1271 |
| 16k vs 8k | ns |
| 32k vs 16k | <0.0001 |
| ***Ae. aegypti*<sup>S</sup> vs *Ae. aegypti*<sup>R</sup>** | **P-value** |
| Blood feeding | ns (0.0798) |
| 4k | ns (0.1048) |
| 8k | 0.0113 |
| 16k | <0.0001 |
| 32k | <0.0001 |

**Fig. S1.**
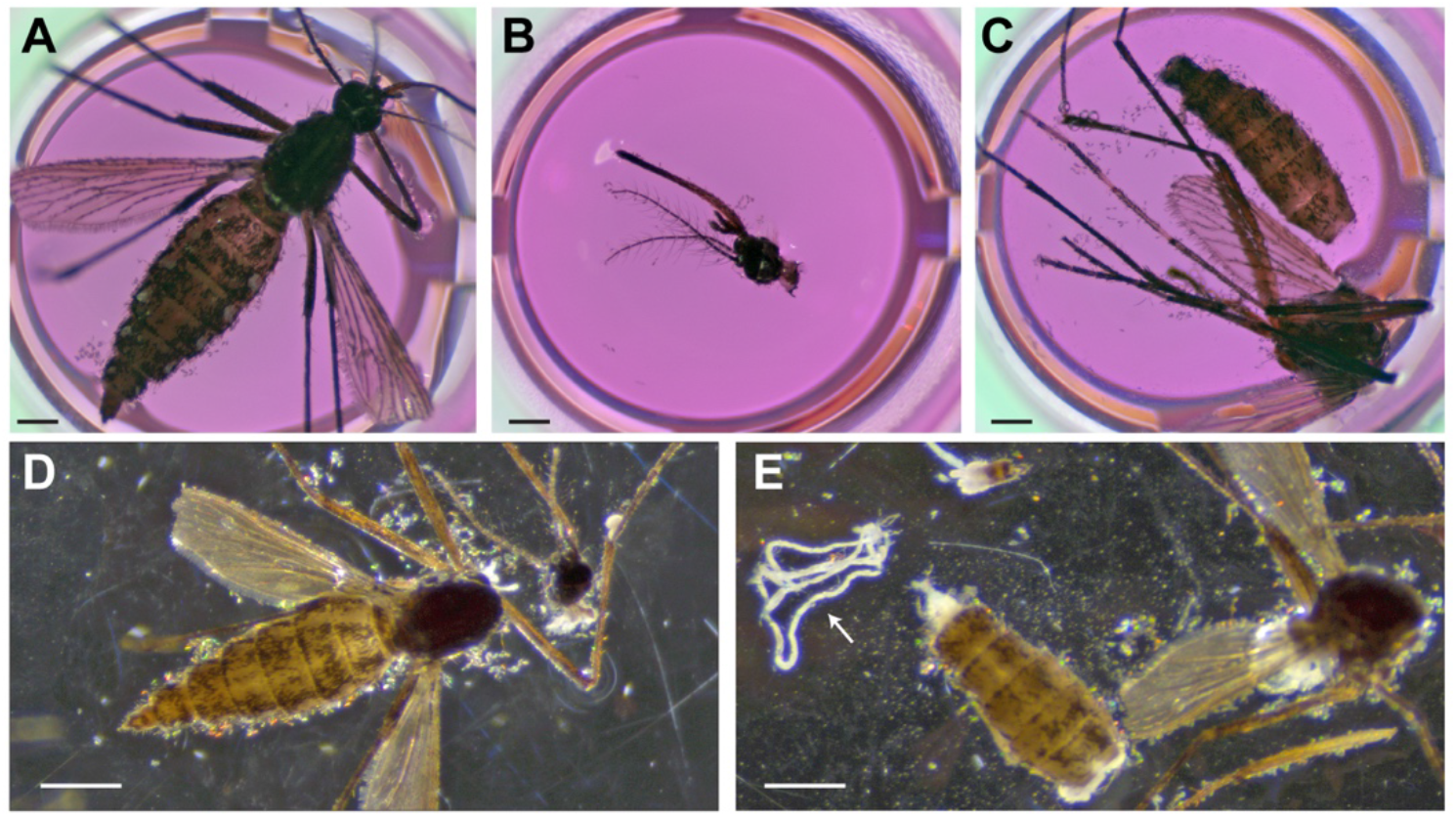
Mosquito dissection for analysis after emergence assay. After the emergence assay from the whole body (**A**), the head (**B**) and carcass (**C**) were placed individually into separate wells. The number of L3 larvae emerging from the dissected head and carcass were assayed after incubation at 37 °C. The Malpighian tubules were removed from the carcass prior to placing it in the well. Mosquito after removing head with fine forceps (**D**) and carcass after the Malpighian tubules (arrow) are dissected out (**E**). The scale bars in A-C and D-E are 500 μm and 1000 μm, respectively.

**Fig. S2.**
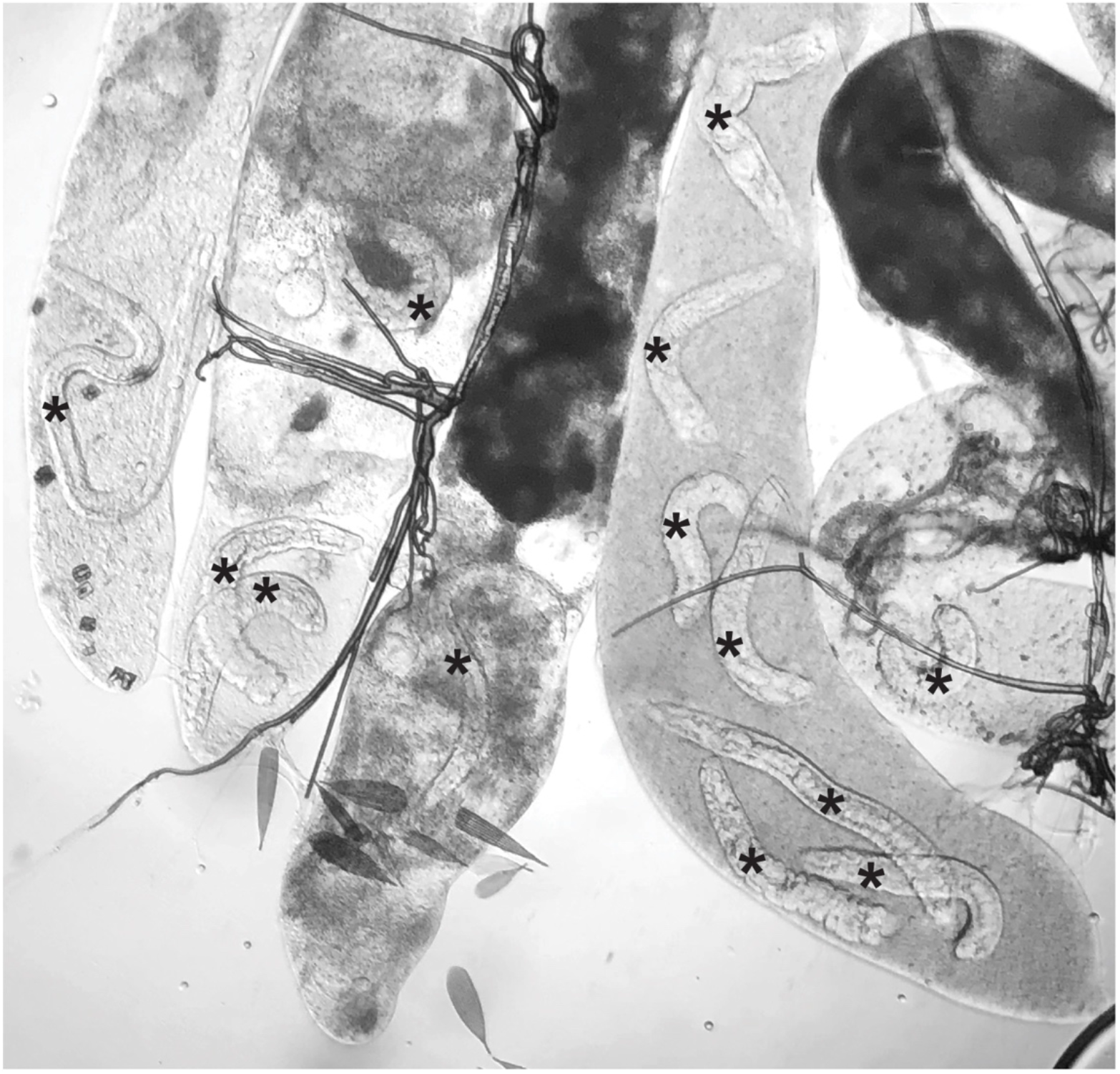
Example of *D. immitis* larval lengths scored in Malpighian tubules. Malpighian tubules dissected following an emergence assay contain viable larvae of different lengths indicated with asterisks. Different stage larvae are typically present.

**Fig. S3.**
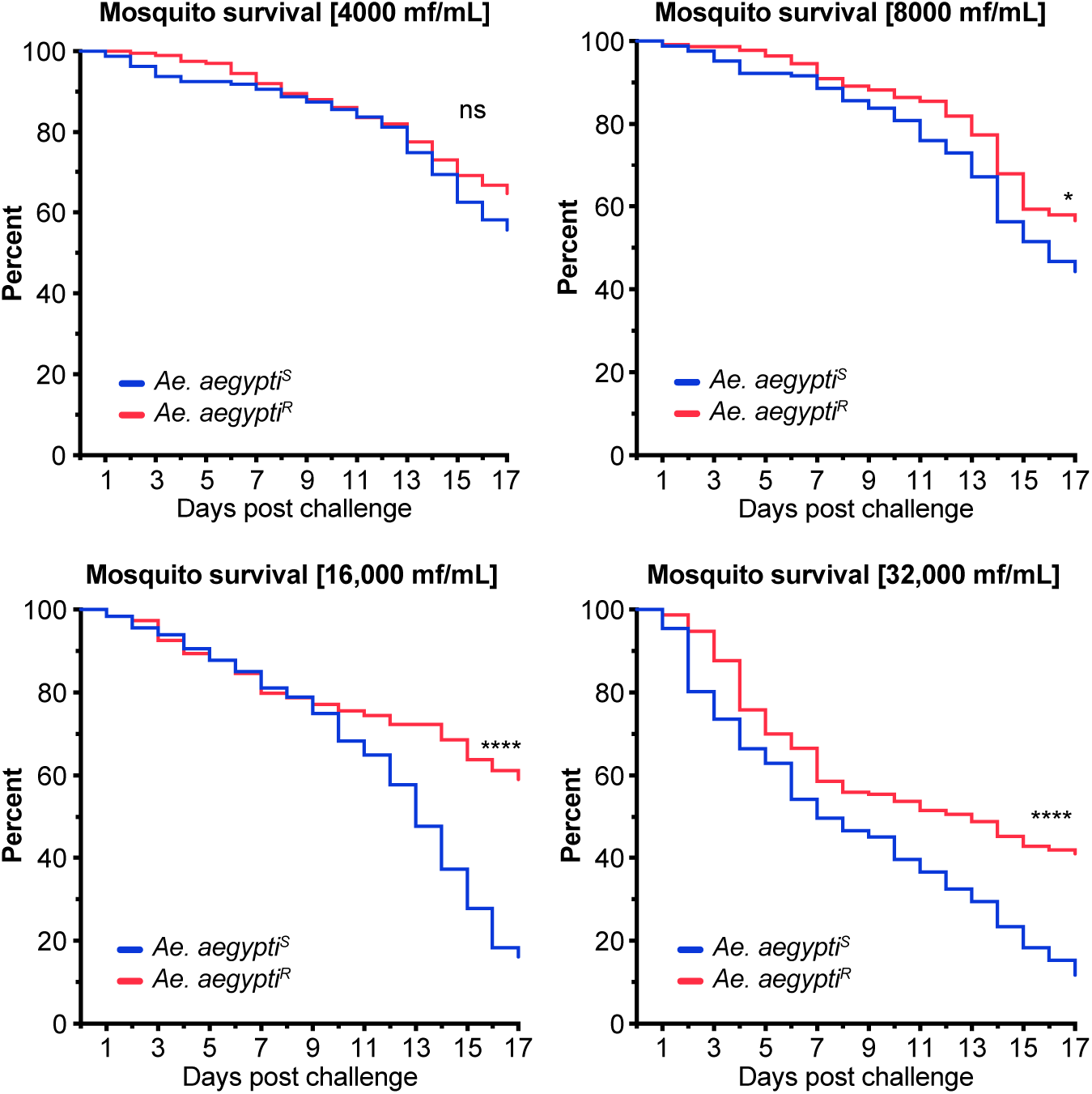
*Ae. aegypti^S^* have greater mortality to than *Ae. aegypti^R^* following *D. immitis* infection. Pairs of adjacent treatment groups from Figure 5D and 5E were analyzed by Kaplan Meier and relationships with significant differences indicated with asterisks here and in Table S1. *Ae. aegypti^S^* and *Ae. aegypti^R^* are indicated in blue and red, respectively.

**Supplemental movie 1.** Two eL3 in a well of a 96-well plate following an emergence assay performed on *Ae. aegypti^S^* 14 days after infection with *D. immitis.* Another eL3 is visible at the top of the frame.

**Supplemental movie 2.** Third-stage larvae visible moving in the proboscis of *Ae. aegypti^S^* 14 days after infection with *D. immitis*.

